# Rapid masking of saccadic motion results from low-velocity input and is largely invariant to movement amplitude

**DOI:** 10.64898/2026.04.03.716410

**Authors:** Wiebke Nörenberg, Richard Schweitzer, Martin Rolfs

**Affiliations:** Department of Psychology, Humboldt-Universität zu Berlin, Germany; Berlin School of Mind and Brain, Humboldt-Universität zu Berlin, Germany; Centro Interdipartimentale di Mente e Cervello, Università degli studi di Trento, Italy; Cluster of Excellence ‘Science of Intelligence’, Technische Universität Berlin, Germany

## Abstract

Saccadic eye movements sweep the visual scene across the retina, yet the resulting motion is rarely perceived. Visual factors alone, such as the presence of static pre– and post-saccadic input, can attenuate motion perception, suggesting effective masking of the motion signal during early visual processing. Here, we isolated the visual component of this reduction in motion perception using simulated saccades presented to fixating observers. Across two experiments, we manipulated motion amplitude (6–18 dva), duration, and velocity profile and measured perceived amplitude and velocity at varying masking durations. Visual masking strongly reduced perceived motion amplitude and velocity, with short halftimes (∼15 ms) that were largely invariant across saccade amplitudes. Critically, motion following a naturalistic saccadic velocity profile was perceived as shorter and slower than constant-velocity motion matched in amplitude and duration, even without explicit masking. This additional reduction increased with both amplitude and duration. These results show that visual mechanisms alone can account for substantial motion reduction across a wide range of amplitudes and demonstrate a partially separable contribution of the saccadic velocity profile, suggesting that the temporal structure of retinal motion itself supports perceptual continuity across eye movements.

## Introduction

Every time we move our eyes, the retinal image is rapidly displaced, yet we remain unaware of both the resulting motion and the displacement of the scene. Here, we focus on the former, referred to as saccadic omission: a lack of conscious experience of the visual consequences of saccades, despite the visual scene travelling across the retina with speeds of several hundred degrees per second (Bridgeman et al., 1994; Campbell & Wurtz, 1978). This perceptual continuity is remarkable given the scale and frequency of saccadic eye movements in everyday vision.

Extra-retinal signals are thought to attenuate visual sensitivity around the time of saccades, a mechanism termed saccadic suppression (Burr et al., 1982; Diamond et al., 2000; see Wurtz, 2018 for a review). However, intra-saccadic input can reach awareness when a stimulus is stabilized on the retina or when post-saccadic visual input is absent (Campbell & Wurtz, 1978; Castet & Masson, 2000; Schweitzer et al., 2025), demonstrating that active saccadic suppression alone is insufficient to account for saccadic omission (see Castet, 2010, for a review of intra-saccadic motion perception). Visual masking by the pre– and post-saccadic scenes has been proposed as a complementary mechanism, and several studies have investigated its contribution to saccadic omission (Ibbotson & Cloherty, 2009). Post-saccadic input appears especially critical: perceived smear during saccades is attenuated when the target remains visible after saccade offset (Bedell & Yang, 2001). The temporal profile of this omission resembles backward masking, where a visual transient impairs the perception of an earlier stimulus within a brief temporal window (B. G. Breitmeyer & Ogmen, 2000). Moreover, transients from spatially distant stimuli additively impair intra-saccadic perception, pointing to contributions from both masking and attentional distraction (Balsdon et al., 2018). Using simulated saccades and high-speed stimulus presentations, which allow visual factors to be isolated from extra-retinal signals, Duyck et al. (2018) demonstrated that visual input alone plays a significant role in attenuating intra-saccadic motion perception, reducing the perceived amplitude of the simulated saccade even in the absence of any eye movement. Replaying the retinal consequences of saccades in a paradigm investigating saccadic motion streaks yields perceptual results virtually identical to those during active saccades (Schweitzer et al., 2025), and visual-only mechanisms activated by image shifts can account for the perceptual properties of saccadic omission without invoking explicit motor commands, with suppressive signals originating as early as the retina (Idrees et al., 2020). Together, these findings establish visual masking as an important component of saccadic omission.

Two questions have nonetheless remained unresolved. First, the only study that used simulated saccades to isolate the role of the visual component to motion attenuation (Duyck et al., 2018) was restricted to a single amplitude of 6 degrees of visual angle (dva), leaving open whether visual masking remains as effective across saccade sizes beyond this amplitude, which would be necessary for this mechanism to contribute to saccadic omission more broadly. This is particularly relevant given evidence that suppression strength scales with saccade amplitude (Stevenson et al., 1986), indicating that mechanisms attenuating the perceptual consequences of saccades do not operate uniformly but adjust dynamically to saccade magnitude. Second, prior work has largely treated the saccade as a uniform motion event, without addressing whether the shape of the saccadic velocity profile, its characteristic acceleration and deceleration (Collewijn et al., 1988), contributes to motion reduction over and above the static scenes flanking the movement; or more generally, which properties of the stimulus, whether its spatial extent, temporal duration, or kinematic structure, determine the strength and completeness of masking. These factors are crucial for understanding the mechanism underlying saccadic omission, because they determine the extent to which visual masking alone can account for it and may further provide the empirical foundation for computational models of motion attenuation during saccadic eye movements.

To address both questions, we simulated the visual consequences of saccades spanning 6 to 18 dva in amplitude and systematically manipulated their duration and velocity profile. We compared a saccadic velocity profile to motion of equivalent amplitude and duration but with constant velocity, allowing the contribution of kinematics to be isolated from that of the flanking masks. We used full-field patterns of repetitive pink noise that emulate the high-contrast, full-field masks present in everyday vision without providing salient spatial endpoints that could anchor judgements of scene displacement and thereby bias motion reports. Across two experiments, we found that visual masking robustly reduces perceived motion amplitude and speed across a wide range of naturalistic saccade amplitudes, and that the saccadic velocity profile contributes additional attenuation even in the absence of explicit masking intervals. We additionally examined whether observers had metacognitive access to masking-induced perceptual degradation by collecting confidence ratings alongside motion reports and found no consistent evidence that confidence tracked the accuracy of perceived motion. These findings demonstrate that purely visual factors robustly attenuate intrasaccadic motion perception, and identify the velocity profile as a contributor to the masking signal that has so far received little attention, with potential relevance for how the visual system maintains perceptual continuity across eye movements.

## Methods

### Participants

Participants were recruited through the “Psychologischer Experimental-Server Adlershof” (PESA) of the Humboldt-Universität zu Berlin. Visual acuity was assessed prior to the first session using a Snellen chart to ensure normal or corrected-to-normal vision (i.e., 20/20 ft. acuity). All participants provided written informed consent before taking part and were compensated with 10€ per hour or course credit. The study was approved by the Ethics Committee of the Department of Psychology at Humboldt-Universität zu Berlin (reference number 2018-36, “Mechanismen visueller Stabilität”) and conducted in accordance with the Declaration of Helsinki (2013).

*Experiment 1.* Fifteen participants were recruited, of whom twelve completed data collection (10 female; mean age ± SD: 24.0 ± 3.6 years; 6 right-eye dominant). One additional participant was excluded as an outlier because their perceived velocity estimates were substantially higher than those of all other participants across conditions (see Supplement 1), leading to model non-convergence, leaving eleven participants in the reported analyses. The preregistration can be found at: https://osf.io/st7du.

*Experiment 2.* Eleven participants were recruited. One participant withdrew from participation, r ten participants in the reported analyses (5 female; mean age ± SD: 26.4 ± 3.5 years; 6 right-eye dominant). The preregistration can be found at https://osf.io/rf5eb.

### Apparatus

Participants were seated 340 cm from a large projection screen (250.2 × 141.0 cm; Stewart Silver 5D Deluxe, Stewart Filmscreen, Torrance, CA) in a dimly lit room with their heads stabilized on a chin rest. Visual stimuli were projected onto the screen via a PROPixx DLP projector (VPixx Technologies, Saint-Bruno, QC, Canada) at a resolution of 960 × 540 pixels and a refresh rate of 1440 Hz. Eye movements were tracked binocularly using a TRACKPixx3 tabletop system (VPixx Technologies, Saint-Bruno, QC, Canada) at a sampling rate of 2000 Hz. Stimulus presentation and response collection were controlled by a Dell Precision T7810 workstation running MATLAB 2018b (MathWorks, Natick, MA, USA) with PsychToolbox (Brainard, 1997; Kleiner et al., 2007). In Experiment 1, responses were given by moving a standard mouse leftward or rightward and confirming by left-clicking. In Experiment 2, participants responded using the left and right arrow keys on a standard keyboard, confirming their reports by pressing the space bar.

### Stimuli

*Noise background.* On each trial, a random repetitive 1/f pink noise pattern (RMS contrast of 0.1664) spanning the full projection screen was generated. This stimulus was chosen for its naturalistic spatial frequency spectrum (Geisler, 2008). The pattern was horizontally periodic, repeating seamlessly every 6 dva to ensure the absence of salient vertical boundaries (see Fig. 1A). The amplitude was matched to the smallest motion amplitude ensuring that the pattern on the screen was identical before and after movement and consistent across all conditions, ensuring that motion percepts were driven by global displacement rather than the tracking of individual features (Duyck et al., 2018).

**Figure 1:**
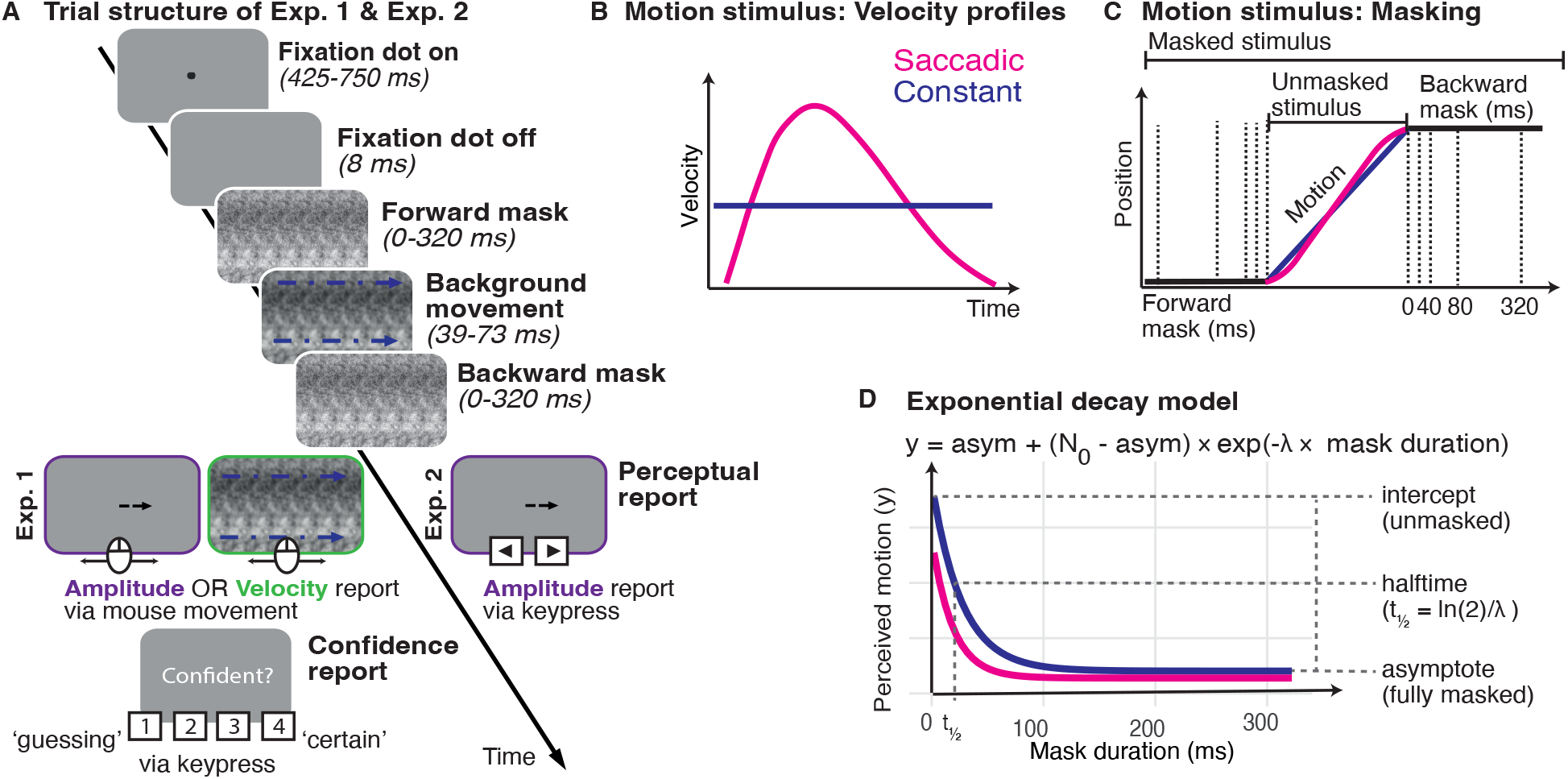
Experimental procedure, stimulus and modeling approach. (A) Trial structure. Each trial began with a fixation period, after which a 1/f pink noise background moved horizontally across the screen. Depending on the masking condition, the motion was preceded by a forward mask and followed by a backward mask of equal duration (0, 20, 40, 80, or 320 ms). Participants subsequently reported the perceived direction and amplitude of the motion using a response arrow. (B) The two velocity profiles used: a naturalistic saccadic profile derived from averaged real saccade recordings (green), characterised by rapid acceleration and deceleration, and a constant velocity profile of equivalent amplitude and motion duration (red). (C) Schematic of the background position over time during a trial. The noise background remained stationary during the forward mask, moved during the motion interval, and returned to a stationary position during the backward mask. Dotted vertical lines indicate the mask duration levels used. The motion trajectory is shown for both the saccadic (pink) and constant velocity (blues) profiles, which were matched in amplitude and motion duration. (D) Schematic of the exponential decay model fitted to perceived amplitude and velocity as a function of mask duration.

*Saccadic velocity profile.* To approximate the kinematics of a natural saccade, we derived an average velocity profile from an existing eye movement dataset (https://osf.io/d2dyu/, Rolfs et al., 2025) that was scaled to match each condition (for details see Supplement 2). Recorded saccades of approximately 12 dva were identified and their velocity profiles were averaged across trials. Constant motion conditions used a constant velocity profile of equivalent amplitude and duration, with no acceleration or deceleration (Fig. 1B).

### Procedure

*General trial structure.* Both experiments followed the same basic trial structure (Fig. 1A). At the start of each trial, a fixation point appeared at the center of the screen. After a fixation control period of 400 ms the trial sequence started. The fixation dot disappeared after an additional period of 25 – 350 ms, leaving the screen empty for 8 ms. Then, the noise background moved either leftward or rightward for a motion duration determined by the experimental condition (39.6, 55.6, or 72.9 ms). Depending on the masking condition, the motion sequence was either unmasked or preceded by a forward mask and followed by a backward mask of equal duration (Fig. 1C). Mask duration had five levels: 0, 20, 40, 80, and 320 ms, where 0 ms indicates no mask was presented. Immediately after the motion sequence, participants were prompted to provide a perceptual report as described below. Trials were repeated at the end of the block if fixation was broken (if an eye position outside of the fixation radius of 1.5 dva around the target was registered), or saccades were detected. Saccades were detected using the Engbert-Kliegl algorithm based on eye velocity calculated from eye position data (Engbert & Kliegl, 2003). Saccades were detected using a velocity threshold of 5 SD, a minimum duration of 30 samples, and merged if separated by fewer than 10 samples(Engbert & Mergenthaler, 2006).

*Experiment 1.* The experiment followed a 2 (velocity profile: constant, saccadic) × 3 (amplitude: 6, 12, 18 dva) × 5 (mask duration: 0, 20, 40, 80, 320 ms) × 2 (direction: left, right) × 10 (repetitions) design, with report type (amplitude or velocity) as a between-session factor, yielding 1200 trials per participant across two sessions of 600 trials each. Critically, amplitude and motion duration covaried according to the main sequence of natural saccades (Bahill et al., 1975), the three amplitude levels were paired with motion durations of 39.6, 55.6, and 72.9 ms respectively, such that stimulus speed increased with amplitude.

In the amplitude report session, participants adjusted the length and direction of a response arrow to match their perceived motion using the mouse, followed by a confidence rating on a 4-point scale (1 = guessing, 2 = unsure, 3 = somewhat confident, 4 = very confident). To encourage participants to judge their own performance rather than task difficulty, they were specifically instructed to choose 4 if they thought their deviation from the presented physical stimulus was minimal (below 10%), 3 if it could have been slightly larger (10–20%), 2 if their report may be significantly different from the true value (above 20%) and 1 if they were guessing and have no confidence in their report.

In the velocity report session, participants adjusted the speed and direction of a moving reference stimulus presented for the average stimulus duration (including masks and motion) of 239.4 ms. To prevent the reference stimulus from masking the motion stimulus, we presented a blank screen for 1 s before the response period started. Reference stimuli alternated with blank screens of 500 ms duration (to prevent motion adaptation and aliasing) until participants confirmed their response with a mouse click, again followed by a confidence rating. They were able to reset the velocity to zero by right-clicking.

*Experiment 2.* The experiment followed a 2 (velocity profile: constant, saccadic) × 3 (amplitude: 6, 12, 18 dva) × 3 (motion durations: 39.6, 55.6, 72.9 ms) × 5 (mask duration: 0, 20, 40, 80, 320 ms) × 2 (direction: left, right) × 12 (repetitions) design, yielding 2160 trials across three sessions of 720 trials each completed on separate days. In Experiment 2, amplitude and motion duration were manipulated independently: The three amplitude levels (6, 12, 18 dva) were each combined with three motion duration levels (39.6, 55.6, 72.9 ms), yielding nine amplitude × motion duration combinations per profile. After each motion sequence, participants adjusted a response arrow using the left and right arrow keys to indicate perceived direction and amplitude and confirmed their response by pressing the space bar. Before the main experiment, participants completed a practice block of 20 trials; each participant achieved more than 80% correct direction reports. All conditions were presented in randomized order; no confidence ratings were collected.

### Data analysis

*Model.* We diverged from the preregistered plan of a repeated measures ANOVA with Bonferroni-corrected pairwise comparisons as the model-based approach allows us to characterize the full temporal decay function. Perceived amplitude and velocity were each modelled as an exponential decay function of mask duration (see Fig. 1D):

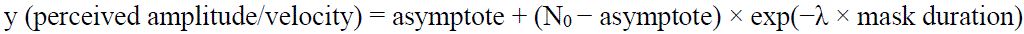

where N_0_, the intercept, is the perceived magnitude in the absence of masking (0 ms mask duration), λ is the decay rate governing how rapidly perception is suppressed with increasing mask duration, and the asymptote is the perceptual floor under full masking. Decay rate is reported as halftime (t½ = ln(2)/λ × 1000 ms), the mask duration at which perceived magnitude reaches the midpoint between N_0_ and the asymptote (Fig. 1D).

*Model fitting.* Models were fit using Bayesian multilevel nonlinear regression implemented in brms (Bürkner, 2021). Weakly informative priors were used for all parameters (Experiment 1, amplitude report: N_0_ ∼ Normal(10, 3), λ ∼ Normal(45, 15), asymptote ∼ Normal(2, 0.7), Experiment 1, velocity report: N_0_ ∼ Normal(60, 30), λ ∼ Normal(45, 15), asymptote ∼ Normal(25, 10), Experiment 2: N_0_ ∼ Normal(6, 3), λ ∼ Normal(50, 20), asymptote ∼ Normal(1, 0.7)). Participant-level random effects were included on all parameters where model complexity permitted, with an LKJ(2) prior on the random effect correlation matrix and Exponential(1) priors on random effect standard deviations. All models were run with 4 chains of 6000 iterations each (warmup = 3000), adapt_delta = 0.995, and max_treedepth = 15. All models converged satisfactorily (max Rhat ≤ 1.006 across all datasets; min bulk ESS ratio ≥ 0.14).

*Model selection.* For each dataset, a sequence of models of increasing parsimony was compared using leave-one-out cross-validation (LOO-IC, for a summary see Supplement 3; Vehtari et al., 2017). Models within one standard error of the best-fitting model were considered equivalent in predictive accuracy, and the most parsimonious among these was selected. Briefly, for Experiment 2 the winning models included a velocity profile × amplitude interaction on N_0_, main effects of both predictors on λ and asymptote, and participant-level intercepts only for the asymptote. For Experiment 1, the winning model additionally included profile × motion duration and amplitude × motion duration interactions on N_0_, and main effects of all three predictors on λ and asymptote.

*Inference.* For Experiment 1, the model intercepts represent the Constant profile at grand mean amplitude. The velocity profile was treatment-coded with Constant as the reference level, such that the profile coefficient directly reflects the Saccadic versus Constant contrast. Amplitude was sum-coded with 12 dva as the implicit level, such that all explicit coefficients reflect deviations from the grand mean within the Constant profile. In Experiment 2, motion duration was additionally sum-coded (with 55.6 ms as the implicit level), and the intercept therefore represents the Constant profile at grand mean amplitude and motion duration. Effects were considered credible if the 95% credible interval (CrI) of the posterior excluded zero; the probability of direction (pd) is reported for all effects as a continuous measure of evidential strength. All analyses were conducted in R/RStudio (RStudio 2024.04.1+748, R version 4.4.0).

*Confidence ratings.* Confidence ratings were collected as a measure of metacognitive access to masking-induced motion reduction. Metacognitive access here refers to the ability of observers to accurately monitor the reliability of their own perceptual reports: if participants have insight into when their motion percepts are degraded by masking, confidence should decrease as masking increases and should correlate negatively with response error, defined as deviation from physical stimulus values. Alternatively, confidence may track self-consistency, defined as the stability of a participant’s responses relative to their own mean, which would reflect awareness of internal response variability rather than of perceptual accuracy per se. A third possibility is that confidence tracks response magnitude, independently of accuracy or consistency. To examine these possibilities, we computed three trial-level measures for each participant: response magnitude, defined as the absolute value of each trial’s motion response (amplitude in dva or velocity in dva/s); absolute response error, defined as the absolute difference between each trial’s response and the physical stimulus value (amplitude in dva or velocity in dva/s), and response self-consistency, defined as the absolute deviation of each trial’s response from that participant’s own mean response within the same condition (amplitude × profile × mask duration). Pearson correlations between confidence and each of these measures were then computed across all trials within each participant and tested against zero using one-sample t-tests.

## Results

### Experiment 1

In Experiment 1, we investigated whether motion masking is effective across a large range of saccade amplitudes and examined the contribution of the saccadic velocity profile. As we used stimuli drawn from the saccadic main sequence, amplitude and motion duration were not manipulated independently. Stimuli followed the natural covariation between saccade size and motion duration, such that 6 dva stimuli always had a motion duration of 39.6 ms, 12 dva of 55.6 ms, and 18 dva of 72.9 ms. Effects attributed to amplitude in Experiment 1 therefore reflect the combined influence of these properties, that is, effects of saccade size as it naturally occurs, rather than a pure effect of spatial displacement.

#### Visual masking robustly reduces perceived motion amplitude and velocity across a wide range of saccade sizes

Perceived amplitude and velocity both decreased exponentially with increasing mask duration across all conditions and both velocity profiles (Fig. 2A-B). In the reference condition (grand mean amplitude, Constant profile), motion amplitude perception decayed rapidly with a halftime of 16.9 ms [14.3, 20.2], reduced to a perceived amplitude of 3.3 dva [2.0, 4.6] at the asymptote, well below veridical. Velocity perception decayed at a comparable rate (halftime: 14.5 ms [12.5, 17.3]), reaching a fully masked percept of 28.6 dva/s [19.5, 37.3]. This pattern extended robustly across the full range of tested saccade sizes. Masking rate did not differ credibly across amplitudes within the Constant profile for amplitude reports (6 dva vs grand mean: pd = 86.9%; 18 dva vs grand mean: pd = 69.5%) but for velocity reports, masking was credibly faster at 6 dva (−1.5 ms [−3.3, −0.6], pd = 99.8%) while 18 dva did not differ credibly from the grand mean (pd = 67.2%), suggesting that the temporal dynamics of masking are largely invariant across the naturalistic saccade range for amplitude reports, with a modest but credible tendency toward faster masking at the smallest amplitude in velocity reports (Fig 2E-F).

**Figure 2.**
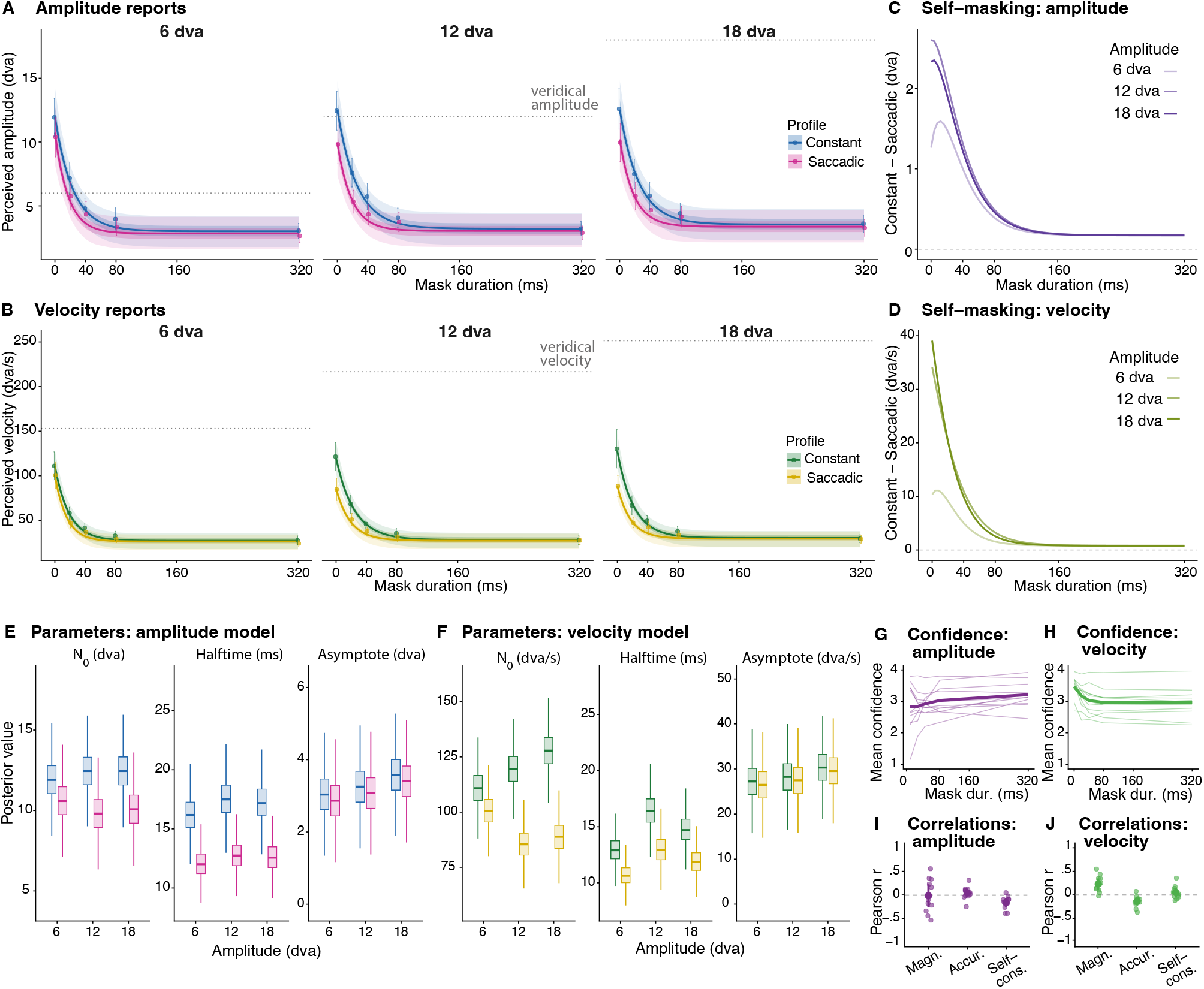
Experiment 1: Effect of static mask durations and velocity profile (self-masking) on perceived amplitude and velocity. A–B) Posterior mean predictions for amplitude reports (A) and velocity reports (B) as a function of mask duration, separately for constant (amplitude: blue, velocity: green) and saccadic (amplitude: pink, velocity: yellow) motion profiles across presented stimulus amplitudes (6, 12, and 18 dva). Shaded bands denote 50% (dark) and 95% (light) credible intervals. Points and error bars show raw condition means ± 1 SE. Dotted horizontal lines indicate physical amplitude (A) and physical mean velocity (B). C–D) Difference curves (Constant minus Saccadic) for amplitude reports (C) and velocity reports (D), showing the magnitude of the velocity profile effect (self-masking of the motion stimulus) as a function of mask duration for all three amplitude conditions, with darker shading indicating larger physical amplitude. E–F) Posterior distributions of the three exponential decay model parameters (N₀, halftime, and asymptote) for the amplitude model (E) and velocity model (F), shown as boxplots per amplitude × profile condition. G–H) Mean confidence ratings from 1 (guessing) to 4 (certain) as a function of mask duration for amplitude reports (G) and velocity reports (H). Thin lines show individual participants; thick lines show the group mean. I–J) Per-participant Pearson correlations between confidence and response magnitude (absolute response magnitude) response accuracy (absolute error relative to physical stimulus) and between confidence and self-consistency (absolute deviation from each participant’s own mean response), for amplitude reports (I) and velocity reports (J). Points show individual participants; central symbols and bars show the group mean ± 95% CI.

Perception of fully masked motion stimuli (corresponding to the asymptote) only weakly increased with amplitude within the Constant profile for both report types (Fig. 2E-F): for amplitude reports, perception for 6 dva stimuli was smaller than the grand mean (−0.25 dva [−0.41, −0.09], pd = 99.9%) while it was increased for 18 dva stimuli (+0.29 dva [+0.13, +0.45], pd = 100%), indicating that stimuli containing larger displacements retain slightly more residual motion percept under full masking. Velocity reports showed a similar pattern (6 dva: −1.36 dva/s [−3.05, +0.32], pd = 94.2%; 18 dva: +1.73 dva/s [+0.05, +3.42], pd = 97.8%), though all conditions remained far below the physical stimulus.

#### The saccadic velocity profile reduces perceived magnitude of unmasked motion, with effects increasing with motion amplitude

Even in the absence of explicit masking intervals, saccadic motion was perceived as smaller than constant at grand mean amplitude (amplitude: −2.10 dva [−2.95, −1.18], pd = 100%; velocity: −27.82 dva/s [−35.88, −19.34], pd = 100%, Fig. 2A-B). Credible profile × amplitude interactions revealed that this attenuation increased with motion amplitude for both report types (Fig. 2E-F). For amplitude reports, the effect was significantly weaker at 6 dva (profile × 6 dva: +0.80 dva [+0.39, +1.21], pd = 100%), indicating reduced attenuation for smaller amplitudes, while the interaction at 18 dva showed an effect in the opposite direction which did not reach credibility (profile × 18 dva: −0.27 dva [−0.68, +0.13], pd = 90.4%). For velocity reports, both interactions were credible and in opposite directions (profile × 6 dva: +17.54 dva/s [+12.95, +22.32], pd = 100%; profile × 18 dva: −11.29 dva/s [−16.11, −6.60], pd = 100%), yielding total profile-related reductions in unmasked velocity perception of −10.28 dva/s at 6 dva, −34.07 dva/s at 12 dva, and −39.11 dva/s at 18 dva (Fig. 2D). The saccadic profile also reduced half times relative to constant velocity motion (amplitude reports: −6.15 ms [–9.68, −3.15], pd = 100%; velocity reports: −3.48 ms [−6.10, −1.22], pd = 99.9%). The difference between constant and saccadic velocity profile conditions was largest without masking (mask duration = 0 ms) and decreased progressively with increasing mask duration, converging toward the fully masked asymptote (Fig. 2C-D). Even though we saw a small reduction in both dependent variables, the saccadic profile did not credibly affect the asymptote for either report type (amplitude reports: −0.18 dva [−0.41, +0.06], pd = 93%; velocity reports: −0.78 dva/s [−3.21, +1.74], pd = 73.3%, Fig. 2E-F).

#### Perception of unmasked motion is far from veridical

Even without masking, perceived amplitude and velocity deviated substantially from the physical stimulus, but in different ways (Fig. 2A-B). The grand mean unmasked perceived amplitude was 12.27 dva [9.70, 14.83], close to the middle stimulus level of 12 dva, but this apparent accuracy masked large individual differences instead of demonstrating accurate perception of the physical stimulus properties: within the Constant profile, unmasked perception at 6 dva fell only modestly below the grand mean (−0.36 dva [−0.69, −0.03], pd = 98.4%) while 18 dva did not differ credibly from it (+0.18 dva [−0.13, +0.50], pd = 86.9%), with condition differences much smaller than inter-individual variability (see also Supplement 1), suggesting that amplitude reports for unmasked motion may reflect stable idiosyncratic priors over motion magnitude rather than accurate encoding of the stimulus. Velocity reports showed a qualitatively different pattern: the grand mean unmasked perceived velocity was 119.48 dva/s [102.81, 136.37], substantially below the range of physical mean speeds (∼153–250 dva/s), with velocity reports systematically underestimating stimulus velocity across all conditions despite credible scaling with motion amplitude (6 dva: −8.55 dva/s [−12.14, −4.96], pd = 100%; 18 dva: +8.45 dva/s [+4.58, +12.47], pd = 100%, Fig. 2E-F).

#### Confidence ratings show no consistent relationship with masking-induced motion reduction

On each trial, participants rated their confidence in their perceptual report on a 4-point scale from ‘guessing’ to ‘certain’. Given that masking substantially reduced perceived motion, we explored whether confidence tracked this degradation, which would be consistent with observers having metacognitive access to the reliability of their motion percepts. Confidence trajectories differed across report types: for amplitude reports, mean confidence slightly increased with mask duration (from 2.84 at 0 ms to 3.21 at 320 ms; Figure 2G), while for velocity reports confidence slightly declined with masking (from 3.47 at 0 ms to 2.96 at 320 ms; Figure 2H). In all cases, confidence remained well above what accurate metacognition would predict given the large deviations from physical amplitude and velocity. Per-participant correlations between confidence and absolute response error were near zero for amplitude reports (mean r = 0.05 [−0.05, 0.14], t(10) = 1.16, p = 0.273; Fig. 2I; for analysis of confidence groups split by response trend see Supplement 4) and weakly negative for velocity reports (mean r = −0.14 [−0.22, −0.05], t(10) = −3.67, p = 0.004; Fig. 2J). For velocity reports, however, the correlation with absolute response magnitude was stronger (mean r = 0.24 [0.12, 0.36], t(10) = 4.48, p = 0.001), indicating that participants were primarily tracking signal strength rather than accuracy. Correlations between confidence and distance from each participant’s own re-sponse mean, a measure for self-consistency, were significantly negative for amplitude reports (mean r = −0.16 [−0.25, −0.06], t(10) = −3.36, p = 0.004), indicating that participants were more confident when their response fell close to their own typical response within that condition, while the equivalent correlation was near zero for velocity reports (mean r = 0.05 [−0.04, 0.15], t(10) = 1.30, p = 0.223). Together, these modality-specific patterns suggest that confidence tracked properties of the response process, signal strength for velocity reports and response typicality for amplitude reports, rather than the accuracy of the underlying motion percept.

### Experiment 2

Experiment 1 established the masking effect across a large range of naturally occurring saccade sizes and the contribution of the saccadic velocity profile using stimuli drawn from the main sequence, in which amplitude and motion duration necessarily covary. Experiment 2 crossed amplitude and motion duration orthogonally across 18 conditions, allowing their independent contributions to be examined by adding 12 conditions which diverge from the main sequence and do not correspond to naturally occurring saccade kinematics. Overall, direction error rates were low but increased significantly with decreased mask duration (for detailed error rate results see Supplement 5).

#### Masking is rapid and robust across all amplitude and motion duration combinations

Perceived amplitude rapidly decreased with increasing mask duration across all 18 conditions (Fig. 3A,D), with a grand mean halftime of 14.3 ms [12.6, 16.5]. Masking was faster for larger amplitudes (18 vs grand mean: −2.3 ms [−3.2, −1.6], pd = 100%; 6 vs grand mean: +1.3 ms [+0.8, +2.1], pd = 100%) and shorter motion durations (39.6 vs grand mean: −2.1 ms [−2.4, −1.7], pd = 100%; 72.9 vs grand mean: +1.6 ms [+1.2, +1.9], pd = 100%). Because amplitude and motion duration have opposing effects on halftime, their contributions partially cancel on the main sequence diagonal, consistent with the intermediate halftimes observed in Experiment 1. The saccadic profile did not credibly affect masking rate (−0.6 ms [−2.1, 0.8], pd = 82.1%, Fig. 3B) across these conditions.

**Figure 3.**
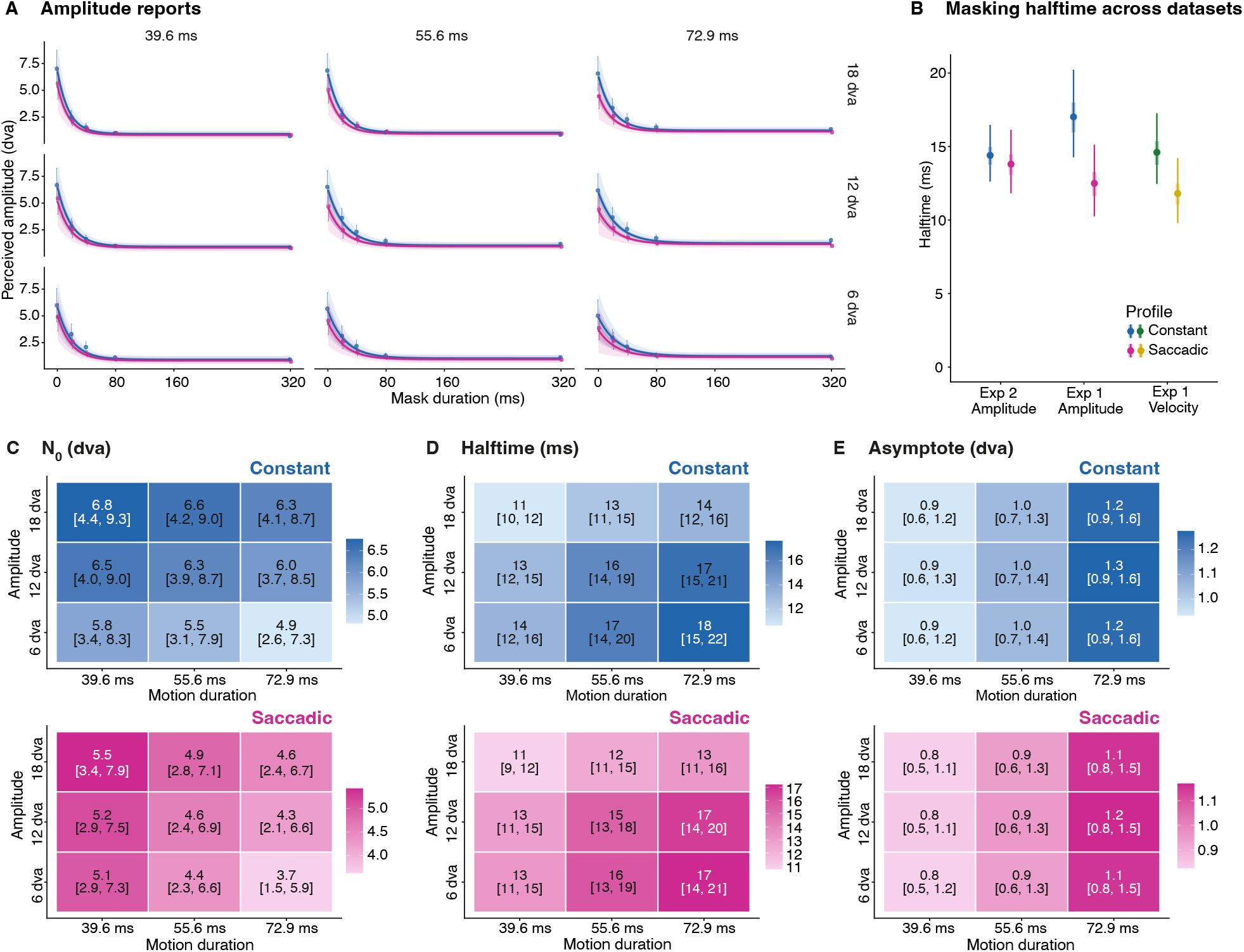
Experiment 2: Effects of stimulus amplitude, motion duration, and motion profile on motion masking. A) Posterior mean predictions for amplitude reports as a function of mask duration, separately for constant (blue) and saccadic (pink) motion profiles, faceted by physical stimulus amplitude (6, 12, and 18 dva) and motion duration (39.6, 55.6, and 72.9 ms). Shaded bands denote 50% (dark) and 95% (light) credible intervals. Points and error bars show observed condition means ±1 SE. B) Masking halftime (in ms) compared across all three datasets (Experiment 2 amplitude reports, Experiment 1 amplitude reports, and Experiment 1 velocity reports), separately for constant and saccadic motion profiles. Points show posterior means; thick and thin bars denote 50% and 95% credible intervals, respectively. Amplitude datasets shown in blue and pink; velocity dataset in green and yellow. C to E) Posterior marginal means of the three time-course model parameters: unmasked perceived amplitude N₀ (C), masking halftime (D), and perceptual floor asymptote (E), displayed as heatmaps over the amplitude × motion duration grid. Each cell shows the posterior mean with 95% credible interval. Top row = Constant profile (blue gradient), bottom row = Saccadic profile (pink gradient); darker fill indicates higher parameter values.

#### The saccadic velocity profile reduces perception of unmasked motion, scaling with both motion amplitude and duration

As in Experiment 1, saccadic motion was perceived as smaller than constant motion without masking (Fig. 3A,C) at grand mean amplitude (−1.39 dva [−1.91, −0.82], pd = 100%). Credible profile × amplitude interactions confirmed that this effect increased with amplitude: profile-driven motion attenuation was reduced at at 6 dva (profile × 6 dva: +0.36 dva [+0.23, +0.50], pd = 100%) and stronger at 18 dva (profile × 18 dva: −0.19 dva [−0.32, −0.05], pd = 99.6%). Independently of amplitude, the profile effect also increased with motion duration (profile × 39.6 ms: +0.25 dva [+0.12, +0.38], pd = 100%; profile × 72.9 ms: −0.17 ms [−0.30, −0.04], pd = 99.4%).

#### The perceptual floor under full masking is determined by motion duration, not amplitude

Fully masked perception in the reference condition (Constant profile, grand mean amplitude and duration) was 1.06 dva [0.74, 1.38]. The parametric manipulation of motion amplitude and duration allowed us to dissociate the motion amplitude effect on the asymptote observed in Experiment 1, revealing that the modest asymptote increase seen there was driven by the covariation of amplitude and motion duration along the main sequence diagonal rather than amplitude per se. Instead, motion duration alone credibly determined the perceptual floor (39.6 ms vs grand mean: −0.15 dva [−0.19, −0.10], pd = 100%; 72.9 ms vs grand mean: +0.18 dva [+0.13, +0.23], pd = 100%) while amplitude had no credible effect (6 dva vs grand mean: −0.01 dva [−0.06, +0.04], pd = 58.7%; 18 dva vs grand mean: +0.00 dva [−0.05, +0.05], pd = 55.0%). Notably, the saccadic profile also credibly reduced the perceptual floor (−0.10 dva [−0.17, −0.02], pd = 99.5%, Fig. 3E), a direction consistent with the trend observed in Experiment 1 that had not reached credibility there (−0.12 dva [−0.36, +0.11], pd = 84.6%, Fig.2 E,F). This indicates that the saccadic velocity profile contributes additional suppression even at the longest mask durations, suggesting that profile-based suppression and mask-based suppression operate at least partially independently rather than converging on the same floor.

#### Masking effects are largely consistent across response modalities and experiments

To verify consistency across experiments more broadly, we refitted the Experiment 1 model structure to the subset of Experiment 2 conditions corresponding to the main sequence diagonal (6 dva / 39.6 ms, 12 dva / 55.6 ms, 18 dva / 72.9 ms). Although absolute parameter estimates differed, reflecting a bias toward larger amplitudes in Experiment 1, the pattern of effects was very consistent across experiments (see Table 1 for full fixed effects across all models). The saccadic profile credibly reduced unmasked perception at grand mean amplitude (−1.60 dva [−1.78, −0.40], pd = 99.7%), with credible profile × amplitude interactions confirming that this attenuation grew with saccade size (profile × 6 dva: +0.51 dva [+0.30, +0.73], pd = 100%; profile × 18 dva: −0.32 dva [−0.54, −0.10], pd = 99.7%). Crucially, the profile also credibly reduced the perceptual floor in this subset (−0.13 dva [−0.24, −0.02], pd = 98.7%), converging with the full Experiment 2 result and supporting the interpretation that the effect is genuine but below the detection threshold of the Experiment 1 data where reports were collected with a potentially less sensitive report modality. The profile effect on halftime was likewise credible in this subset (−2.37 ms [−4.06, −1.09], pd = 99.9%), confirming the corresponding result of Experiment 1 (amplitude reports: pd = 100%, velocity reports: pd = 99.9%) but not the full Experiment 2 model (pd = 82.1%). This suggests that the effect observed in Experiment 1 is genuine, indicating that the saccadic profile accelerates masking specifically when stimuli follow natural saccade kinematics.

**Table 1.** Fixed effects from all Bayesian nonlinear mixed models, mixed contrast coding. Each column corresponds to one model fit. Exp 1 (amp) and Exp 1 (vel) = amplitude and velocity report models from Experiment 1. Exp 2 = full factorial model from Experiment 2; Exp 2 (diagonal) = Experiment 1 model structure refitted to the main-sequence subset of Experiment 2 (6 dva / 39.6 ms, 12 dva / 55.6 ms, 18 dva / 72.9 ms). Motion profile is treatment-coded (reference = Constant); amplitude is sum-coded (deviation from grand mean; 12 dva is the implicit level with deviation = −(b6 + b18)). N_0_ = perceived motion in the absence of static masks. Halftime = mask duration at the midpoint between N_0_ and asymptote (ms); coefficients for halftime predictors derived via delta method. Asym = perceptual floor under full masking. pd = probability of direction. Green shading indicates that the 95% credible interval excludes zero; grey text indicates that it does not.

| Fixed effects of all models |  |  |  |  |  |  |  |  |
| --- | --- | --- | --- | --- | --- | --- | --- | --- |
| Predictor | Exp 1 (amp) |  | Exp 1 (vel) |  | Exp 2 |  | Exp 2 (diagonal) |  |
|  | Estimate [95% CrI] | pd | Estimate [95% CrI] | pd | Estimate [95% CrI] | pd | Estimate [95% CrI] | pd |
| **$N_0$** |  |  |  |  |  |  |  |  |
| Intercept (Constant profile, grand mean amplitude) | 12.27 [9.7, 14.83] | 100% | 119.48 [102.81, 136.37] | 100% | 6.11 [3.75, 8.51] | 100% | 6.45 [4.03, 9.07] | 100% |
| Profile: Saccadic vs Constant | -2.1 [-2.95, -1.18] | 100% | -27.82 [-35.88, -19.34] | 100% | -1.39 [-1.91, -0.82] | 100% | -1.6 [-2.24, -0.94] | 100% |
| Amplitude: 6 dva vs grand mean | -0.36 [-0.69, -0.03] | 98.4% | -8.55 [-12.14, -4.96] | 100% | -0.67 [-0.91, -0.44] | 100% | -0.28 [-0.44, -0.11] | 99.9% |
| Amplitude: 18 dva vs grand mean | 0.18 [-0.13, 0.5] | 86.9% | 8.45 [4.58, 12.47] | 100% | 0.49 [0.25, 0.73] | 100% | 0.12 [-0.05, 0.29] | 91% |
| Duration: 39.2 ms vs grand mean |  |  |  |  | 0.31 [0.07, 0.56] | 99.3% |  |  |
| Duration: 71.6 ms vs grand mean |  |  |  |  | -0.35 [-0.57, -0.12] | 99.8% |  |  |
| Profile × Amplitude (6 dva) | 0.8 [0.39, 1.21] | 100% | 17.54 [12.95, 22.32] | 100% | 0.36 [0.23, 0.5] | 100% | 0.51 [0.3, 0.73] | 100% |
| Profile × Amplitude (18 dva) | -0.27 [-0.68, 0.13] | 90.4% | -11.29 [-16.11, -6.6] | 100% | -0.19 [-0.32, -0.05] | 99.6% | -0.32 [-0.54, -0.1] | 99.7% |
| Profile × Duration (39.2 ms) |  |  |  |  | 0.25 [0.12, 0.38] | 100% |  |  |
| Profile × Duration (71.6 ms) |  |  |  |  | -0.17 [-0.3, -0.04] | 99.4% |  |  |
| Amplitude × Duration (6 × 39.2) |  |  |  |  | 0.12 [0.03, 0.22] | 99.4% |  |  |
| Amplitude × Duration (6 × 71.6) |  |  |  |  | -0.18 [-0.28, -0.09] | 100% |  |  |
| Amplitude × Duration (18 × 39.2) |  |  |  |  | -0.06 [-0.16, 0.03] | 91% |  |  |
| Amplitude × Duration (18 × 71.6) |  |  |  |  | 0.09 [0, 0.19] | 97.9% |  |  |
| **Halftime (ms)** |  |  |  |  |  |  |  |  |
| Intercept (Constant profile, grand mean amplitude) | 16.88 [14.27, 20.23] | 100% | 14.5 [12.46, 17.27] | 100% | 14.33 [12.62, 16.48] | 100% | 16.25 [14.26, 18.55] | 100% |
| Profile: Saccadic vs Constant | -6.15 [-9.68, -3.15] | 100% | -3.48 [-6.1, -1.22] | 99.9% | -0.64 [-2.05, 0.78] | 82.1% | -2.37 [-4.06, -1.09] | 99.9% |
| Amplitude: 6 dva vs grand mean | -0.82 [-2.38, 0.51] | 86.9% | -1.84 [-3.34, -0.55] | 99.8% | 1.34 [0.81, 2.12] | 100% | -0.47 [-1.34, 0.71] | 83.3% |
| Amplitude: 18 dva vs grand mean | 0.27 [-0.87, 1.36] | 69.5% | 0.19 [-0.97, 1.15] | 67.2% | -2.3 [-3.24, -1.57] | 100% | -0.23 [-1.32, 0.58] | 67.8% |
| Duration: 39.2 ms vs grand mean |  |  |  |  | -2.08 [-2.42, -1.74] | 100% |  |  |
| Duration: 71.6 ms vs grand mean |  |  |  |  | 1.55 [1.24, 1.86] | 100% |  |  |
| **Asym** |  |  |  |  |  |  |  |  |
| Intercept (Constant profile, grand mean amplitude) | 3.29 [1.98, 4.55] | 100% | 28.58 [19.53, 37.26] | 100% | 1.06 [0.74, 1.38] | 100% | 1.13 [0.77, 1.52] | 100% |
| Profile: Saccadic vs Constant | -0.18 [-0.41, 0.06] | 93% | -0.78 [-3.21, 1.74] | 73.3% | -0.1 [-0.17, -0.02] | 99.5% | -0.13 [-0.24, -0.02] | 98.7% |
| Amplitude: 6 dva vs grand mean | -0.25 [-0.41, -0.09] | 99.9% | -1.36 [-3.05, 0.32] | 94.2% | -0.01 [-0.06, 0.04] | 58.7% | -0.15 [-0.23, -0.07] | 100% |
| Amplitude: 18 dva vs grand mean | 0.29 [0.13, 0.45] | 100% | 1.73 [0.05, 3.42] | 97.8% | 0 [-0.05, 0.05] | 55% | 0.16 [0.08, 0.24] | 100% |
| Duration: 39.2 ms vs grand mean |  |  |  |  | -0.15 [-0.19, -0.1] | 100% |  |  |
| Duration: 71.6 ms vs grand mean |  |  |  |  | 0.18 [0.13, 0.23] | 100% |  |  |

## Discussion

Saccadic omission has been attributed to a combination of visual masking and extra-retinal suppression, yet their relative contributions have proven difficult to disentangle. By rendering motion of repetitive pink noise patterns at high temporal resolution and manipulating mask duration, motion duration, stimulus amplitude and velocity profile of simulated saccades, we quantified purely visual motion reduction in the absence of eye movements and characterized its temporal dynamics across a large range of saccades sizes. The visual transients resulting from the static masks alone rapidly reduced perceived motion amplitude and velocity across all tested conditions. We further find a motion attenuating effect of the kinematic profile of the simulated movement which is partially separable and occurs even in the absence of static masks. Confidence ratings showed no consistent relationship with masking-induced motion reduction with modality-specific patterns suggesting that confidence tracked properties of the response process rather than the response accuracy, suggesting a dissociation between the motion representation affected by masking and the signals contributing to confidence judgments.

The temporal dynamics of motion masking were strikingly consistent across both experiments and for both amplitude and velocity reports, with halftimes of approximately 15 ms. That masking reduces perceived spatial extent and perceived speed at the same rate, despite these being measured through entirely different response procedures, suggests that both measures are constrained by the same underlying signal. This may reflect attenuation of a common motion signal at an early stage of processing. This interpretation is consistent with motion-energy models, in which speed and displacement are derived from shared spatiotemporal filters with transient impulse responses (Adelson & Bergen, 1985) and with evidence that visual masking operates on the order of tens of milliseconds and disrupts early sensory representations (Breitmeyer & Ganz, 1976; Breitmeyer & Ogmen, 2006). These halftimes are also consistent with recent work establishing that visual masking contributes to saccadic omission (Balsdon et al., 2018; Duyck et al., 2018; Schweitzer et al., 2025), and extend it by showing that pre– and post-stimulus visual transients alone, without any eye movement or extra-retinal signal, are sufficient to reduce perceived motion by approximately 50 % within 15 ms across a parametric range of mask durations and saccade sizes. This complements evidence from paradigms comparing real and replayed saccades suggesting that visual factors contribute substantially to motion reduction during natural eye movements (Schweitzer et al., 2025), though how visual and extra-retinal signals interact during real saccades and their relative weighting under natural viewing conditions remain open questions.

Interestingly, perception of unmasked motion stimuli tracked neither physical amplitude nor velocity veridically: perceived amplitude compressed systematically across a threefold range of physical amplitudes and responded in opposite directions to amplitude and motion duration manipulations which have opposite effects on stimulus speed. While participants preserved relative speed ordering across conditions, absolute velocity reports clustered around 120 dva/s despite average stimulus speeds ranging from 150 to 250 dva/s. This is consistent with reported reductions in speed discrimination at higher velocities (De Bruyn & Orban, 1988). The brief stimulus and report durations may have contributed to this, as short presentation times limit temporal integration and thereby reduce motion discrimination performance (Watamaniuk & Sekuler, 1992). Additionally, pink noise stimuli contain a broad range of spatial frequencies, which may reduce motion signal reliability, effectively producing smearing (D. Burr, 1980), at higher temporal frequencies and thereby impairing velocity estimation (Kelly, 1979).

We show that the simulated saccadic profile contributes to motion reduction over and above the external masks and quantify this contribution. Rather than unifying internal and external sources of motion masking, however, the data suggest they may make at least partially separable contributions to the total magnitude of motion reduction: The saccadic profile not only significantly reduces the perceived motion magnitude of unmasked motion, it also slightly but credibly reduced the perceptual floor (asymptote) in Experiment 2 and in the diagonal refitting of Experiment 2 while both datasets of Experiment 1 exhibited a similar trend. This indicates that profile-based motion attenuation contributes even when external masking has already strongly reduced motion perception. Several explanations for these profile effects are plausible. First, the gradual acceleration and deceleration of the saccadic profile reduces the suddenness of the velocity onset and offset relative to constant velocity motion, which starts at full speed and constitutes a stronger and more abrupt signal for velocity-tuned mechanisms. Thus, the saccadic profile may initially produce a weaker motion signal, or may even act as a masking signal itself below a certain velocity threshold. Second, the velocity changes within the saccadic profile itself may generate competing motion signals corresponding to the acceleration and deceleration phases, interfering with perception of the motion as a coherent whole. Third, motion featuring the saccadic velocity profile reaches higher peak velocities which would increase the above-mentioned motion smearing and may thereby render the signal easier to mask. These explanations are not mutually exclusive, and the present design cannot discern between them. Larger, faster saccades may therefore feature stronger motion attenuation by the visual component alone, not because a stronger extra-retinal signal is generated, but because the eye movement itself generates stronger masking via its temporal profile. Whether this contributes to the effectiveness of perceptual continuity across the naturally occurring range of saccade sizes is speculative, but it is consistent with the known scaling of saccade kinematics, where peak velocity increases systematically with amplitude and larger saccades exhibit broader temporal profiles including longer acceleration and deceleration phases (Bahill et al., 1975; Van Opstal & Van Gisbergen, 1987). Given that motion-sensitive mechanisms respond strongly to spatiotemporal structure (Adelson & Bergen, 1985), these kinematic differences may influence the strength of motion-related responses at early visual encoding stages. However, in the present design the same averaged velocity profile was scaled across amplitudes, which limits how directly these results speak to natural saccade kinematics and means that the observed scaling of masking strength may partly reflect properties of the imposed profile rather than naturalistic variation in saccade dynamics. Future work using empirically recorded saccade trajectories would be required to fully dissociate these factors.

Residual motion perception does survive under full masking, increasing with motion duration rather than amplitude, as demonstrated by the dissociation between motion duration and amplitude effects on the fully masked motion percept in Experiment 2. This is consistent with temporal integration accumulating motion evidence that masking cannot fully eliminate. However, this residual signal remains a small fraction of what accurate encoding would predict.

The pattern of effects was largely consistent across both experiments, perceptual variables, response modalities, and model structures, supporting the interpretation that the reported effects reflect properties of the visual system rather than procedural or model-specific artefacts. Attenuation of unmasked motion perception by the saccadic velocity profile replicated in direction, in its interaction with saccade amplitude, and in approximate magnitude across all four analyses. The opposing effects of amplitude and motion duration on halftime, and their partial cancellation along the main sequence diagonal, likewise replicated across experiments and account for the intermediate halftimes observed in Experiment 1. Two effects showed inconsistent credibility across experiments and perceptual reports. The profile effect on the fully masked percept did not reach credibility in the Experiment 1 amplitude or velocity datasets but was credible in both Experiment 2 and the diagonal refitting; the consistent direction across all datasets supports the interpretation that this effect is small but present. The profile effect on half times was credible in Experiment 1 and the diagonal refitting but not in the full Experiment 2 model. One explanation is that the profile accelerates masking specifically when stimuli follow naturalistic saccade kinematics (Rolfs et al., 2025), such that the effect is attenuated in the full orthogonal design by the inclusion of off-diagonal amplitude-motion duration combinations that do not correspond to natural saccades; whether this reflects a genuine kinematic specificity or a dependence on the parameter combinations sampled remains an open question. However, converging evidence that perception of high-speed motion is lawfully related to saccade kinematics comes from a recent study (Rolfs et al., 2025b), which showed that the visibility of a small moving object during fixation scales with the main sequence, with visibility thresholds scaling proportional to saccadic peak velocity. Notably, using a saccadic velocity profile increased rather than decreased object visibility relative to constant velocity motion. Although the stimuli and tasks differ substantially from those used here, our results complement this finding by showing that for whole-field motion the same kinematic structure instead reduces perceived motion. This opposing pattern may reflect the low-velocity content of the saccadic profile: the gradual acceleration and deceleration phases that may help detect a moving object against a static, uniform background are likely contributing to masking of whole-field motion. A further difference between experiments is that absolute amplitude reports were notably larger in Experiment 1 than Experiment 2, with intercepts of 12.27 and 6.11 dva respectively. The two experiments used different response modalities: in Experiment 1, participants adjusted an on-screen arrow by moving a mouse, a method that is fast but likely less precise, whereas in Experiment 2 participants held down an arrow key until the desired amplitude was reached, a slower method that affords finer perceptual discrimination and more deliberate responding. We therefore attribute the higher amplitude reports in Experiment 1 to response imprecision rather than a genuine difference in perceived amplitude. Critically, halftime estimates were highly comparable across all datasets (14.33-16.88 ms), indicating that the dynamics of masking were unaffected by the response modality. This suggests that while the two methods may differ in their sensitivity to absolute amplitude, they converge on the same relative reduction in motion perception with decreasing mask duration, and report magnitudes should be interpreted in relative rather than absolute terms.

Participants showed no clear evidence of metacognitive access to masking-induced motion reduction. Although confidence trajectories differed across report types, neither followed the pattern expected if observers were sensitive to the perceptual attenuation produced by masking: a steep decline of confidence as masking reduces motion perception. Instead, confidence remained largely decoupled from the magnitude of perceptual change. This dissociation is supported by trial-level analyses. For velocity reports, confidence was only weakly related to response accuracy but showed a stronger correlation with response magnitude, indicating that participants primarily tracked the subjective strength of the motion signal rather than the accuracy of their perceptual report. For amplitude reports, confidence was unrelated to response accuracy or magnitude and instead correlated with self-consistency meaning participants feel most confident when reporting closest to their own typical response within a condition (Caziot & Mamassian, 2021). Rather than reflecting genuine insight into perceptual reliability, confidence appears to have tracked features of the response process, specifically signal strength and self-consistency. The substantial reductions in perceived amplitude and velocity induced by masking were not accompanied by corresponding reductions in confidence. This dissociation is consistent with motion masking disrupting early motion representations (B. G. Breitmeyer & Ogmen, 2000), while confidence and perceptual decisions draw on partially dissociable evidence accumulation processes (Balsdon et al., 2020), leaving confidence insensitive to changes in the fidelity of the underlying motion representation. Taken together, these findings suggest that masking acts primarily on perceptual encoding, producing large changes in the sensory representation of motion, while confidence is computed from a partially distinct decision-level representation that does not encode these changes.

Together, these findings provide further evidence that purely visual transients are sufficient to produce rapid and robust attenuation of perceived motion across a large range of saccade sizes, with consistent temporal dynamics of approximately 15 ms across conditions, experiments, and response modalities. This was demonstrated using broadband full-field stimuli whose spatial frequency content approximates natural visual scenes, suggesting that the reported effects are likely to generalize beyond laboratory conditions. Beyond external masking, we quantify for the first time the self-masking properties of the saccadic velocity profile itself, showing that the kinematic structure of the movement contributes independently to motion attenuation with effects that may persist even when external masking has strongly reduced perceived motion.

## Supplements

### Supplement 1: Individual participant model fits for perceived amplitude and velocity reports

**Figure S1:**
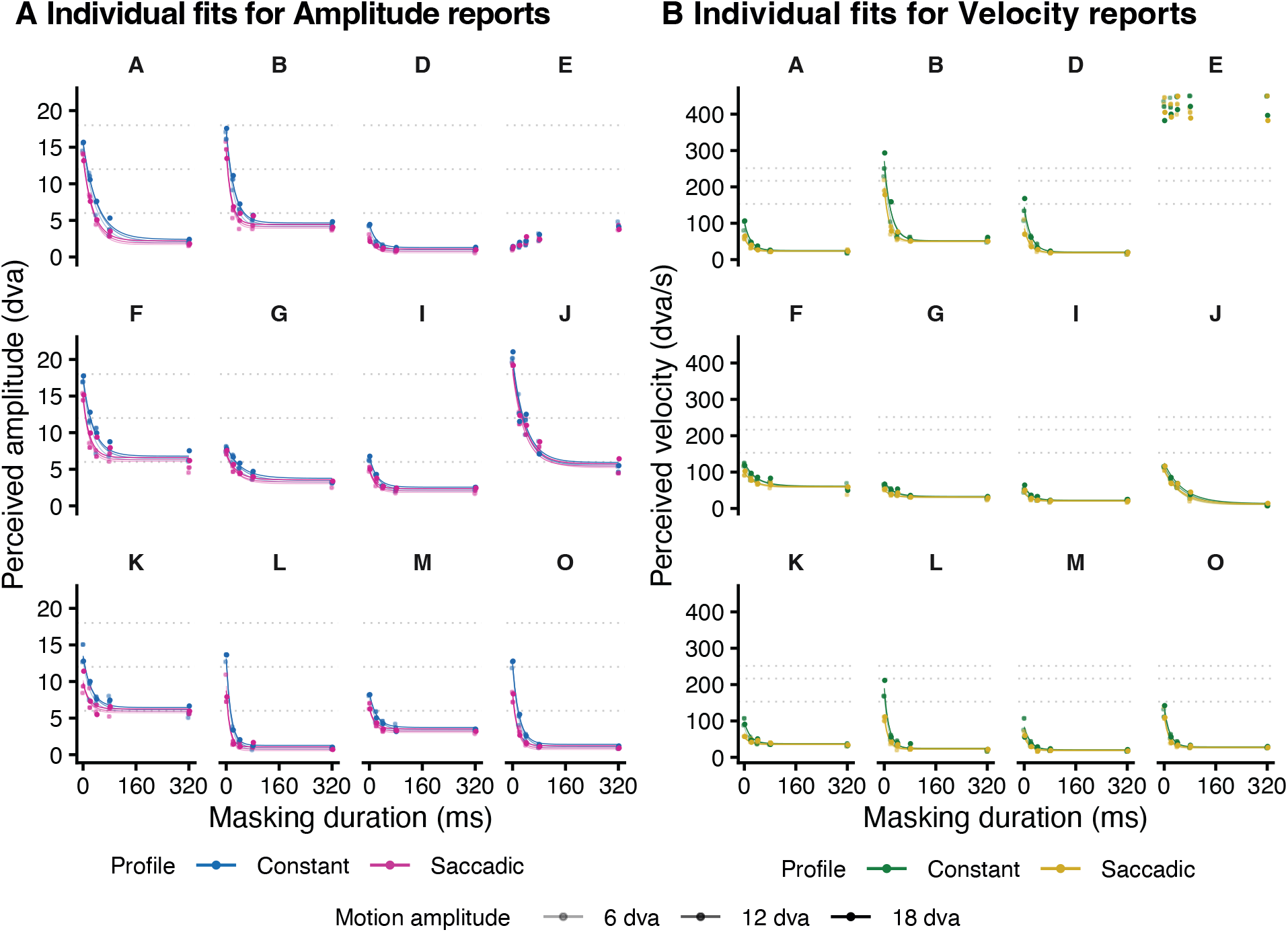
Individual participant data and model fits for perceived amplitude and velocity reports for Experiment 1. **A)** Fitted model curves overlaid on averaged raw data data points for perceived amplitude (dva) as a function of mask duration (ms) for individual participants. Dotted lines indicate physical amplitudes. **B)** Corresponding fits and averaged raw data for perceived velocity (dva/s). Dotted lines indicate physical mean motion velocities. In both panels, blue/green curves represent the Constant motion profile and pink/yellow curves the Saccadic motion profile. Shade indicates motion amplitude (6, 12, and 18 dva). Participant E was excluded from group analyses due to extreme velocity reports and model non-convergence; data shown here for completeness.

### Supplement 2: Velocity profiles of all motion stimuli used in Experiments 1 and 2

**Figure S2.**
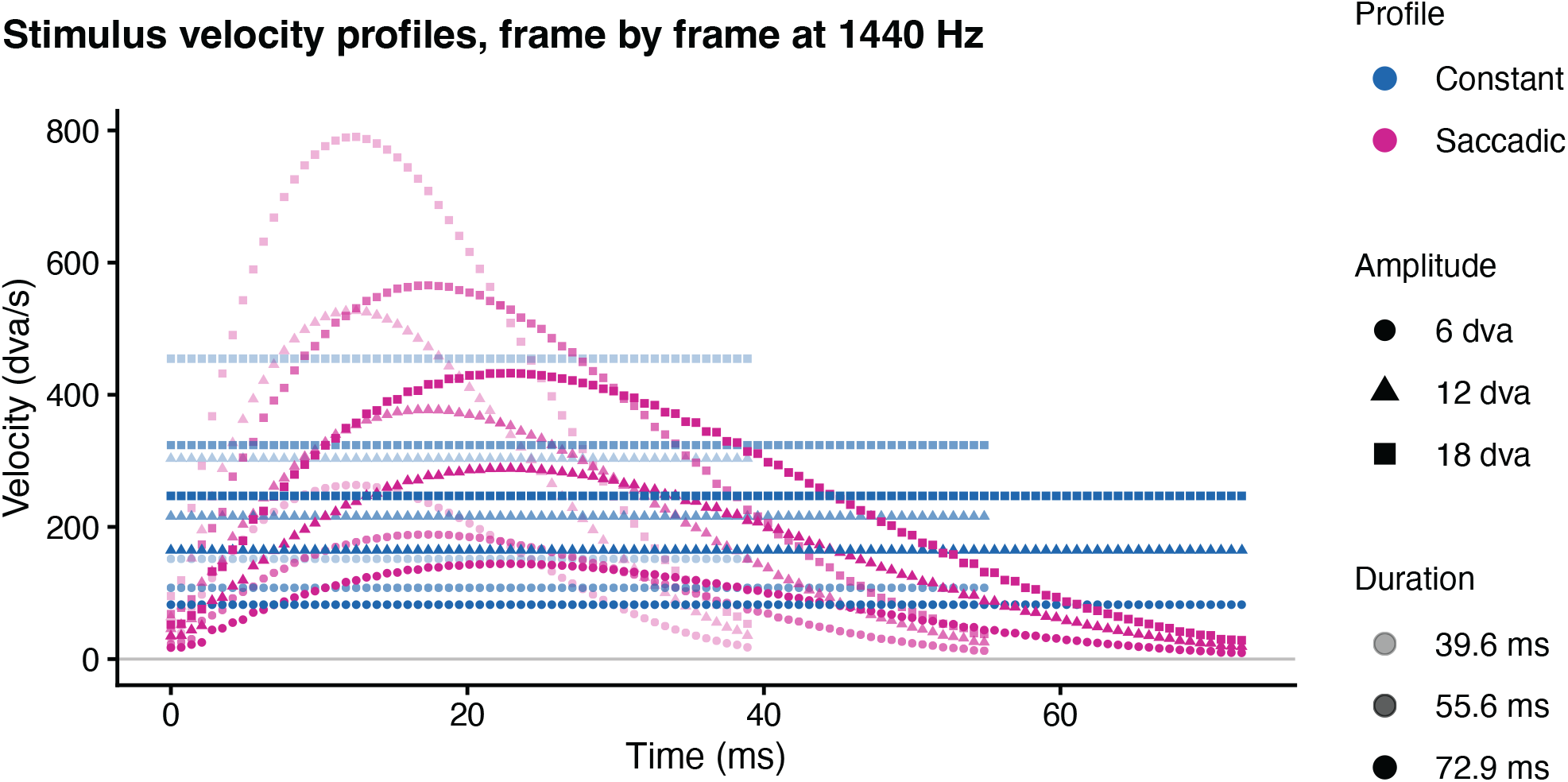
Visualization of velocity profiles. Velocity over time for each of the 18 motion conditions (3 amplitudes × 3 motion durations × 2 profiles), plotted frame-by-frame at the 1440 Hz refresh rate of the display. Constant-velocity profiles (blue) had a velocity of *v* = amplitude / motion duration throughout the motion interval. Saccadic profiles (pink) were obtained by scaling the normalised average position trajectory of recorded ∼12 dva saccades (Rolfs et al., 2025) by the desired amplitude in dva and resampling to the desired duration in ms, then differentiating with respect to time. Symbol shape encodes amplitude (circle = 6 dva, triangle = 12 dva, square = 18 dva) and symbol opacity encodes motion duration (faintest = 39.6 ms, medium = 55.6 ms, fully opaque = 72.9 ms).

### Supplement 3: Bayesian model selection via leave-one-out cross-validation

**Supplementary Table S1.**
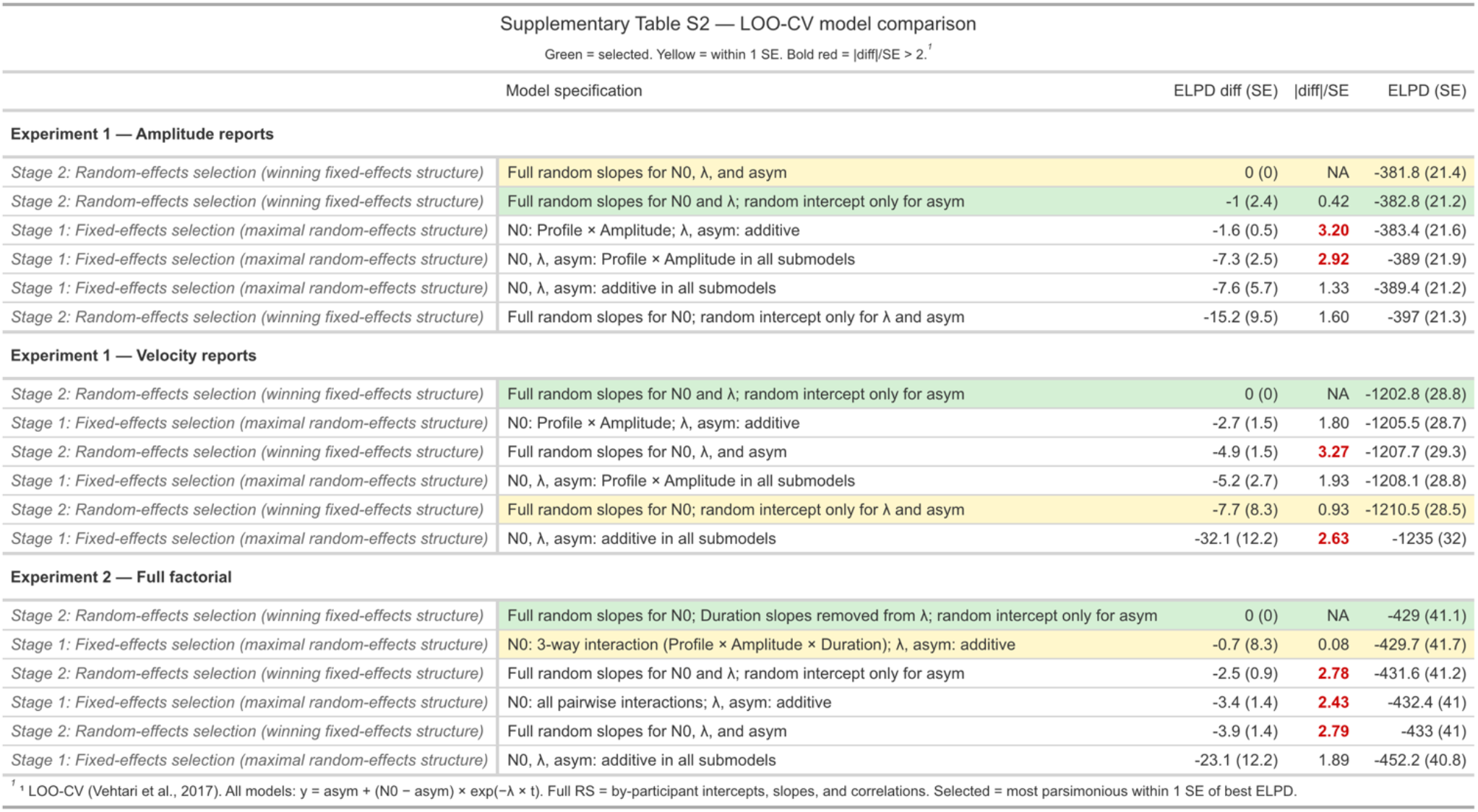
LOO-CV model comparison across all three analyses. Models were selected in two stages: Stage 1 varied the fixed-effects specification of the unmasked perceived motion parameter N_0_ while holding the random-effects structure maximal; Stage 2 held the winning fixed-effects structure and varied the random-effects structure. The selected model (green) is the most parsimonious model within 1 SE of the best ELPD. Yellow shading indicates models within 1 SE of the best. Bold red |diff|/SE values indicate a reliable difference from the best model. ELPD diff = difference in expected log pointwise predictive density relative to the best model; SE = standard error of this difference; |diff|/SE = ratio used to assess reliability of the difference. N_0_ = perceived motion in the absence of masking intervals; λ = decay rate; asym = perceptual floor under full masking. Full random slopes = by-participant intercepts, slopes, and their correlations. LOO-CV computed with moment matching where applicable (Vehtari et al., 2017).

### Supplement 4: Confidence ratings and metacognitive access to masking

The main results establish that confidence tracked response-process properties rather than perceptual accuracy across both report types. Here we characterize the sources of individual variability in this relationship. Participants split roughly evenly on the sign of their amplitude magnitude–confidence correlation (5 positive, 6 negative; Figure S3A). The negative-correlation group reported substantially higher confidence at 0 ms than the positive-correlation group for amplitude reports (M = 3.58 vs 2.42, t(7.3) = 4.37, p = .003), converging at 320 ms (p = .500; Fig. S3B, top). For velocity reports, the positive-correlation group also tended toward higher confidence at both 0 ms (M = 3.69 vs 3.29, t(7.0) = −2.27, p = .057) and 320 ms (M = 3.25 vs 2.72, t(8.3) = −2.08, p = .070; Fig. S3B, bottom), though neither reached significance. For amplitude reports, the negative-correlation group had both a lower unmasked percept (N0: 7.53 vs 15.46 dva, t(8.7) = −4.70, p = .001) and a lower perceptual floor under full masking (asymptote: 1.95 vs 4.90 dva, t(6.4) = −3.15, p = .018; Fig. S3C, top), while masking rate did not differ between groups (halftime: 15.3 vs 19.6 ms, p = .361). For velocity reports, we saw a similar trend, but none of the decay parameters differed credibly between groups (all p > .24; Fig. S3C, bottom). The fact that amplitude perception differed between groups at both end points but masking dynamics did not, and that velocity perception showed no credible group differences, indicates that the group split captured differences in decision-level response style rather than differential sensitivity to masking. Whether the bias in amplitude responses reflects genuine perceptual sensitivity or response strategy cannot be determined from the present data. However, this result converges with the self-consistency result (Fig. S2D) on the same conclusion: confidence tracked decision-level variables rather than masking-induced reductions in motion perception.

**Figure S3.**
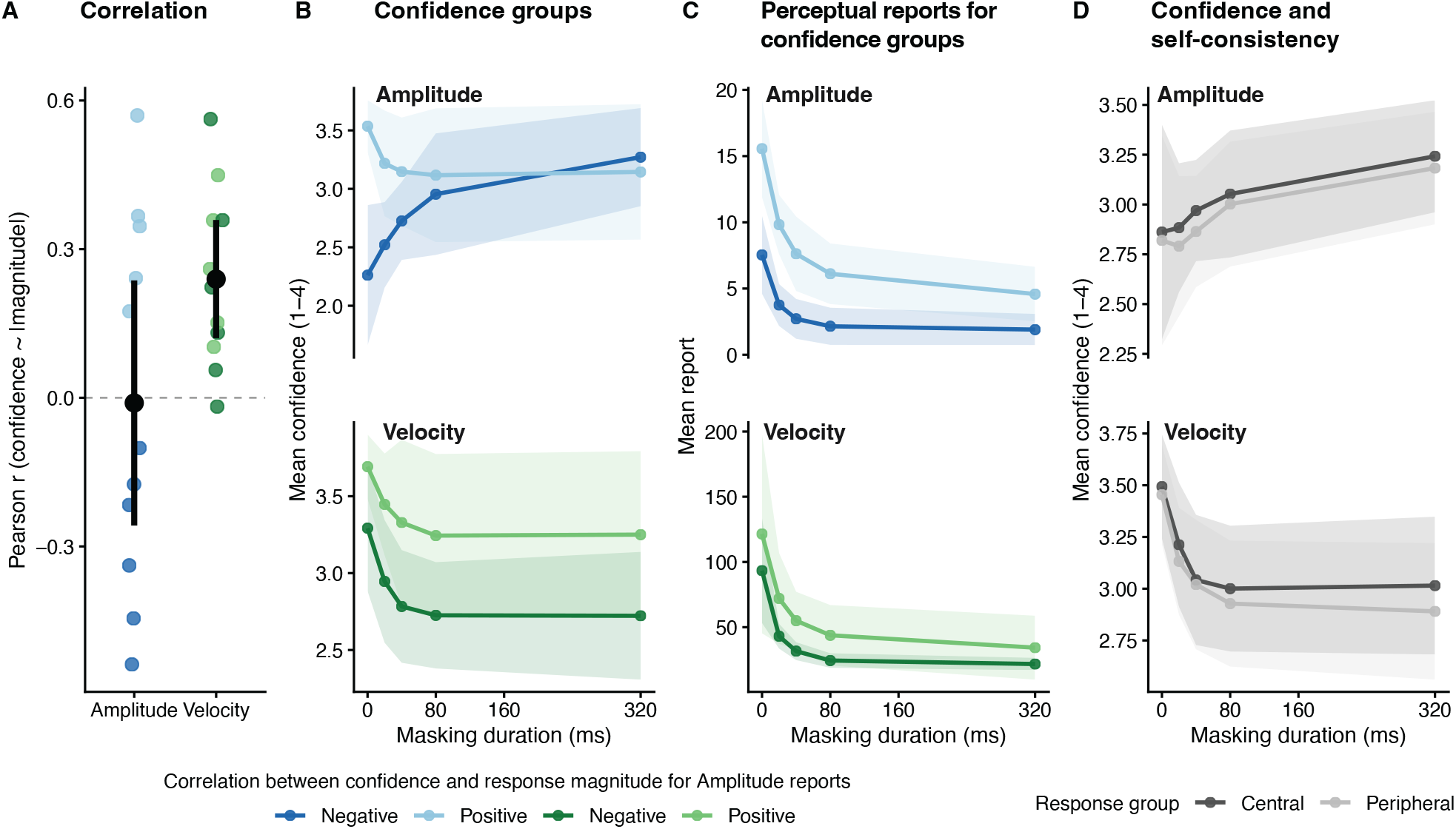
Individual differences in confidence and metacognitive sensitivity. (A) Per-participant Pearson correlations between confidence ratings and response magnitude for amplitude (blue shades) and velocity (green shades) reports. Points show individual participants; central symbols and bars show group mean ± 95% CI. Color indicates the sign of the magnitude–confidence correlation, used as a grouping variable in panels B and C. (B) Mean confidence as a function of mask duration for positive-r and negative-r groups, separately for amplitude (top) and velocity (bottom) reports. Groups are defined by the amplitude-derived r-sign. Shaded bands show ± 95% CI. (C) Mean perceived amplitude (top) and velocity (bottom) as a function of mask duration for the same groups as in B, illustrating that the negative-correlation group had lower perceived motion at both endpoints for amplitude reports but not for velocity reports, while masking dynamics were equivalent between groups in both sessions. Shaded bands show ± 95% CI. (D) Mean confidence as a function of mask duration for trials falling within the central 50% versus outer 50% of each participant’s response distribution within each mask duration level, separately for amplitude (top) and velocity (bottom) reports, illustrating the self-consistency effect reported in the main results. Shaded bands show ± 95% CI.

### Supplement 5: Direction Error Rates

Direction error rates were overall low but decreased monotonically with increasing mask duration (M = 0.08, 0.07, 0.06, 0.03, and 0.01 for 0, 20, 40, 80, and 320 ms respectively, see Fig. S4). A one-way repeated-measures ANOVA confirmed a significant main effect of mask duration (F(4, 36) = 5.33, p = 0.002; Greenhouse-Geisser corrected: p = 0.028), and a paired t-test (data collapsed across conditions) confirmed significantly higher error rates at 0 ms than at 320 ms (mean difference = 0.07, t(9) = 3.62, p = 0.006, 95% CI [0.027, 0.115]). Error rates showed substantial between-participant variability (individual means: 0.004–0.20). Inspection of individual conditions revealed that elevated error rates for unmasked stimuli were concentrated in conditions with the longest amplitude (18 dva), particularly at the shortest motion duration (39.6 ms), which features the highest mean velocity (451 dva/s). This suggests that directional encoding failures reflect difficulty resolving stimulus direction at highest speeds rather than a general consequence of masking.

**Figure S4.**
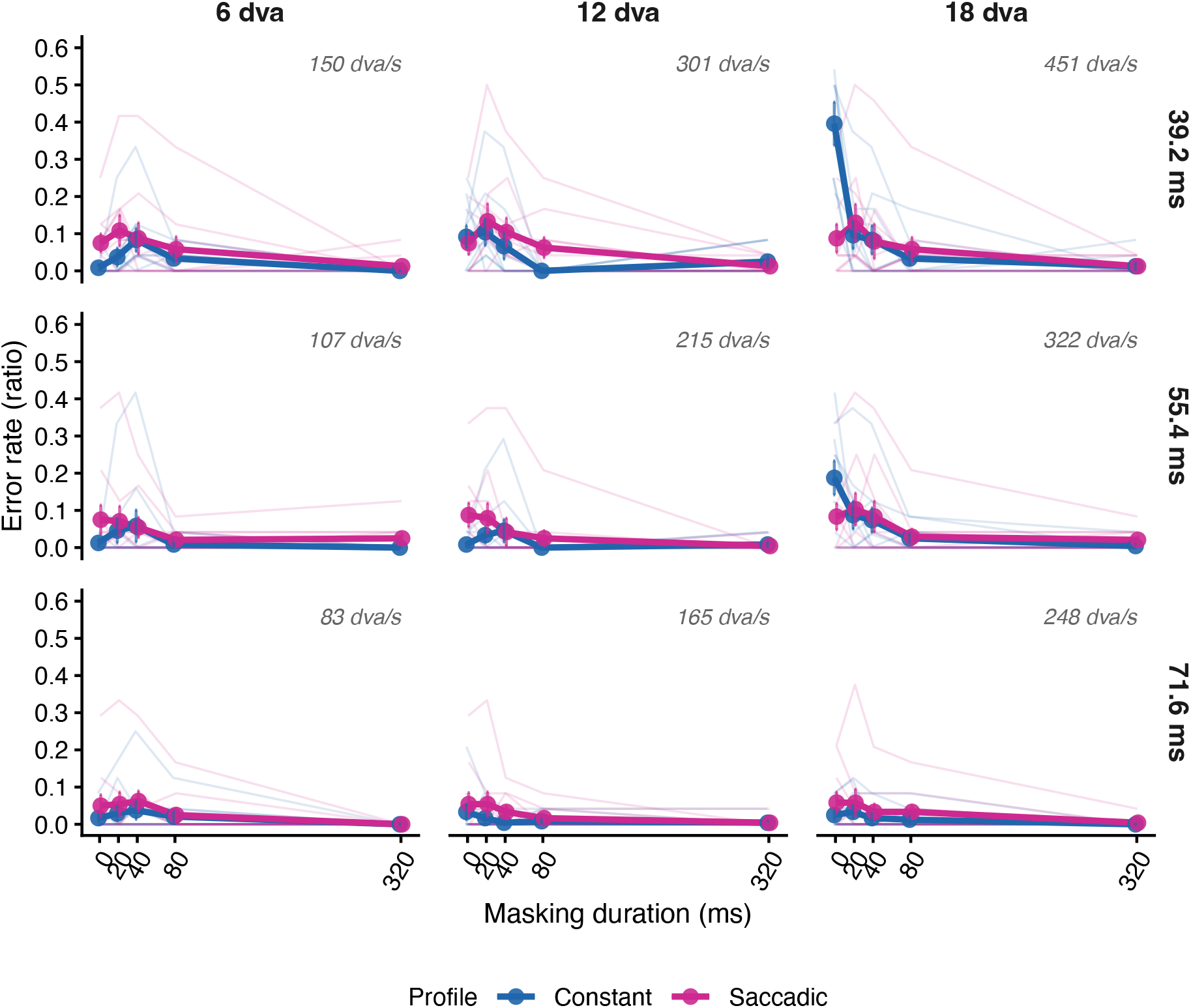
Direction error rates as a function of mask duration in Experiment 2. Each panel shows error rates for constant (blue) and saccadic (pink) motion profiles as a function of mask duration, for all amplitude × motion duration conditions sorted by mean velocity of the motion stimulus (in the upper right corner of each panel). Group mean error rates are shown as thick lines with points; error bars denote ±1 SE. Faint lines show individual participants.

## Acknowledgements

We thank Thomas Symank, Antonia Keller and Mara Doering for their support with data collection and piloting of the experiments, and members of the Laboratoire des Systemes Perceptifs in Paris for comments and suggestions regarding the analysis of confidence reports.

Supported by the European Research Council (ERC) under the European Union’s Horizon 2020 research and innovation programme (grant agreement No. [865715 – VIS-A-VIS]) and Deutsche Forschungsgemeinschaft (DFG) under Germany’s Excellence Strategy – EXC 2002/1 “Science of Intelligence” – project number 390523135.

## References

1. Adelson, E. H., & Bergen, J. R. (1985). Spatiotemporal energy models for the perception of motion. Journal of the Optical Society of America A, 2(2), 284. 10.1364/JOSAA.2.000284

2. Anderson, S. J., & Burr, D. C. (1987). Receptive field size of human motion detection units. Vision Research, 27(4), 621–635. 10.1016/0042-6989(87)90047-2

3. Bahill, A. T., Clark, M. R., & Stark, L. (1975). The main sequence, a tool for studying human eye movements. Mathematical Biosciences, 24(3–4), Article 3–4. 10.1016/0025-5564(75)90075-9

4. Balsdon, T., Schweitzer, R., Watson, T. L., & Rolfs, M. (2018). All is not lost: Post-saccadic contributions to the perceptual omission of intra-saccadic streaks. Consciousness and Cognition, 64, 19–31. 10.1016/j.concog.2018.05.004

5. Balsdon, T., Wyart, V., & Mamassian, P. (2020). Confidence controls perceptual evidence accumulation. Nature Communications, 11(1), Article 1. 10.1038/s41467-020-15561-w

6. Bedell, H. E., & Yang, J. (2001). The attenuation of perceived image smear during saccades. Vision Research, 41(4), 521–528. 10.1016/S0042-6989(00)00266-2

7. Brainard, D. H. (1997). The Psychophysics Toolbox. Spatial Vision, 10(4), 433–436. 10.1163/156856897X00357

8. Breitmeyer, B. G., & Ganz, L. (1976). Implications of sustained and transient channels for theories of visual pattern masking, saccadic suppression, and information processing. Psychological Review, 83(1), 1–36. 10.1037/0033-295X.83.1.1

9. Breitmeyer, B. G., & Ogmen, H. (2000). Recent models and findings in visual backward masking: A comparison, review, and update. Perception & Psychophysics, 62(8), 1572–1595. 10.3758/BF03212157

10. Breitmeyer, B., & Ogmen, H. (2006). Visual Masking. Oxford University Press. 10.1093/acprof:oso/9780198530671.001.0001

11. Bridgeman, B., Van Der Heijden, A. H. C., & Velichkovsky, B. M. (1994). A theory of visual stability across saccadic eye movements. Behavioral and Brain Sciences, 17(2), 247–258. 10.1017/S0140525X00034361

12. Bürkner, P.-C. (2021). Bayesian Item Response Modeling in *R* with **brms** and *Stan*. Journal of Statistical Software, 100(5). 10.18637/jss.v100.i05

13. Burr, D. (1980). Motion smear. Nature, 284(5752), 164–165. 10.1038/284164a0

14. Burr, D. C., Holt, J., Johnstone, J. R., & Ross, J. (1982). Selective depression of motion sensitivity during saccades. The Journal of Physiology, 333(1), 1–15. 10.1113/jphysiol.1982.sp014434

15. Burr, D. C., Morrone, M. C., & Ross, J. (1994). Selective suppression of the magnocellular visual pathway during saccadic eye movements. Nature, 371(6497), Article 6497. 10.1038/371511a0

16. Campbell, F. W., & Wurtz, R. H. (1978). Saccadic omission: Why we do not see a grey-out during a saccadic eye movement. Vision Research, 18(10), Article 10. 10.1016/0042-6989(78)90219-5

17. Castet, E. (2010). Intrasaccadic motion perception. *Dynamics of Visual Motion Processing: Neuronal*, Behavioral and Computational Approaches, 213–238.

18. Castet, E., & Masson, G. S. (2000). Motion perception during saccadic eye movements. Nature Neuroscience, 3(2), Article 2. 10.1038/72124

19. Caziot, B., & Mamassian, P. (2021). Perceptual confidence judgments reflect self-consistency. Journal of Vision, 21(12), Article 12. 10.1167/jov.21.12.8

20. Collewijn, H., Erkelens, C. J., & Steinman, R. M. (1988). Binocular co-ordination of human vertical saccadic eye movements. The Journal of Physiology, 404(1), 183–197. 10.1113/jphysiol.1988.sp017285

21. De Bruyn, B., & Orban, G. A. (1988). Human velocity and direction discrimination measured with random dot patterns. Vision Research, 28(12), 1323–1335. 10.1016/0042-6989(88)90064-8

22. Diamond, M. R., Ross, J., & Morrone, M. C. (2000). Extraretinal Control of Saccadic Suppression. The Journal of Neuroscience, 20(9), 3449–3455. 10.1523/JNEUROSCI.20-09-03449.2000

23. Duyck, M., Wexler, M., Castet, E., & Collins, T. (2018). Motion Masking by Stationary Objects: A Study of Simulated Saccades. I-Perception, 9(3), Article 3. 10.1177/2041669518773111

24. Engbert, R., & Kliegl, R. (2003). Microsaccades uncover the orientation of covert attention. Vision Research, 43(9), Article 9. 10.1016/S0042-6989(03)00084-1

25. Engbert, R., & Mergenthaler, K. (2006). Microsaccades are triggered by low retinal image slip. Proceedings of the National Academy of Sciences, 103(18), Article 18. 10.1073/pnas.0509557103

26. Geisler, W. S. (2008). Visual Perception and the Statistical Properties of Natural Scenes. Annual Review of Psychology, 59(1), Article 1. 10.1146/annurev.psych.58.110405.085632

27. Ibbotson, M. R., & Cloherty, S. L. (2009). Visual Perception: Saccadic Omission — Suppression or Temporal Masking? Current Biology, 19(12), R493–R496. 10.1016/j.cub.2009.05.010

28. Idrees, S., Baumann, M. P., Franke, F., Münch, T. A., & Hafed, Z. M. (2020). Perceptual saccadic suppression starts in the retina. Nature Communications, 11(1), 1977. 10.1038/s41467-020-15890-w

29. Kelly, D. H. (1979). Motion and vision II Stabilized spatio-temporal threshold surface. Journal of the Optical Society of America, 69(10), 1340. 10.1364/JOSA.69.001340

30. Kleiner, M., Brainard, D., & Pelli, D. (2007). What’s new in Psychtoolbox-3?

31. Rolfs, M., Schweitzer, R., Castet, E., Watson, T. L., & Ohl, S. (2025a). Lawful kinematics link eye movements to the limits of high-speed perception. Nature Communications, 16(1), 3962. 10.1038/s41467-025-58659-9

32. Rolfs, M., Schweitzer, R., Castet, E., Watson, T. L., & Ohl, S. (2025b). Lawful kinematics link eye movements to the limits of high-speed perception. Nature Communications, 16(1), 3962. 10.1038/s41467-025-58659-9

33. Schweitzer, R., Doering, M., Seel, T., Raisch, J., & Rolfs, M. (2025). Saccadic omission revisited: What saccade-induced smear looks like. Psychological Review. 10.1037/rev0000574

34. Van Opstal, A. J., & Van Gisbergen, J. A. M. (1987). Skewness of saccadic velocity profiles: A unifying parameter for normal and slow saccades. Vision Research, 27(5), 731–745. 10.1016/0042-6989(87)90071-X

35. Vehtari, A., Gelman, A., & Gabry, J. (2017). Practical Bayesian model evaluation using leave-one-out cross-validation and WAIC. Statistics and Computing, 27(5), 1413–1432. 10.1007/s11222-016-9696-4

36. Watamaniuk, S. N. J., & Sekuler, R. (1992). Temporal and spatial integration in dynamic random-dot stimuli. Vision Research, 32(12), 2341–2347. 10.1016/0042-6989(92)90097-3

37. Wurtz, R. H. (2018). Corollary Discharge Contributions to Perceptual Continuity Across Saccades. Annual Review of Vision Science, 4(1), 215–237. 10.1146/annurev-vision-102016-061207

